# Stage-Specific NF-κB RelA and IRF4 Programs Drive α-Synuclein–Induced Disease-Associated Microglia Differentiation

**DOI:** 10.64898/2026.09.25.754514

**Authors:** Ya-Ting Yang, Yves Greatti, Aidan T. Miller, Jhodi M. Webster, Asta Jurkuvenaite, Josh Randolph, Osama Al-Dalahmah, David A. Figge, Ashley S. Harms

**Affiliations:** Department of Neurology, Killion Center for Neurodegeneration and Experimental Therapeutics, University of Alabama at Birmingham, Birmingham, AL; Aligning Science Across Parkinson’s (ASAP) Collaborative Research Network, Chevy Chase, MD, 20815; Department of Pathology & Cell Biology, Columbia University Irving Medical Center, New York, NY, USA; Taub Institute for Research on Alzheimer’s Disease and the Aging Brain, Columbia University Irving Medical Center, New York, NY, USA; Department of Pathology, University of Alabama at Birmingham, Birmingham, Alabama 35294

## Abstract

Microglial activation in response to aggregated *α*-synuclein (*α*-syn) accumulation is a pathological hallmark of Parkinson’s disease (PD). A distinct activated microglial state, disease-associated microglia (DAM), has been identified across multiple neurodegenerative disorders, including PD. However, the transcriptional and epigenetic programs governing DAM differentiation remain unclear. Here, we show that neuronal *α*-syn overexpression drives progressive microglial state transitions toward DAM, accompanied by increased phagocytic activity and enhanced cytokine and chemokine production. Integrative transcriptomic and chromatin accessibility analyses reveal a hierarchical NF-κB–AP-1–IRF regulatory program underlying these state transitions. Microglial NF-κB p65 (RelA) or IRF4 deficiency impairs microglial progression toward DAM, with RelA and IRF4 governing early microglial activation and subsequent DAM differentiation, respectively. Enrichment of NF-κB, AP-1, and IRF motifs in DAM in human PD supports conservation of this regulatory program across species. Together, our findings establish RelA and IRF4 as sequential, stage-specific regulators that coordinate *α*-syn-induced DAM differentiation in PD.

## Introduction

Parkinson’s Disease (PD) is characterized by progressive loss of dopaminergic neurons in the substantia nigra pars compacta (SNpc) and the accumulation of aggregated *α*-synuclein (*α*-syn) in Lewy bodies and Lewy neurites^1,2^. As the primary brain-resident immune cells, microglia continuously survey the brain microenvironment and respond to *α*-syn aggregation and neuronal degeneration ^3–5^. *In vivo* imaging and postmortem studies demonstrate sustained microglial activation near *α*-syn pathology across disease stages^6,7^, including increased expression of major histocompatibility complex class II (MHCII)^8,9^, suggesting that *α*-syn induces a persistent inflammatory state in microglia. Together these observations implicate microglial activation as a key driver of chronic neuroinflammation and disease progression in PD.

Consistent with this persistent activation, disease-associated microglia (DAM) emerge as a distinct transcriptional state across multiple neurodegenerative disorders, including PD^10,11^. The DAM state is characterized by the downregulation of homeostatic genes and upregulation of programs involved in phagocytosis, lysosomal processing, lipid metabolism, antigen presentation, and chemokine/cytokine signaling^12,13^. DAM have been identified in human and multiple mouse models of neurodegenerations, where they accumulate near pathological lesions and increase during disease progression^14–17^, suggesting that they represent a conserved microglial response to persistent neurodegenerative cues. Despite their association with disease, it remains unclear whether these transcriptional changes translate to microglial function. Moreover, the mechanisms by which persistent neurodegenerative signals establish and maintain the DAM transcriptional state remain poorly understood.

Microglial activation has been implicated as a prerequisite for DAM differentiation^18,19^, yet how sustained activation signals are integrated through transcriptional and epigenetic programs to drive microglial state transitions toward DAM remains unclear. In PD, *α*-syn activates toll-like receptors (TLRs), including TLR2 and TLR4, leading to nuclear factor kappa B (NF-κB) activation^20,21^. During sustained macrophage activation, NF-κB cooperates with other transcription factors (TFs) to establish long-term epigenetic changes and drive a faster or stronger effector response upon subsequent stimulation^22,23^. However, whether sustained activation signals such as *α*-syn induce permanent microglial state transitions toward DAM in an NF-κB-dependent manner remains a critical gap in understanding microglial fate during neurodegeneration.

Here, we investigated how sustained microglial activation driven by neuronal *α*-syn governs microglial state transitions toward DAM using an adeno-associated viral (AAV) mouse model of PD. We identify a stepwise, overlapping enrichment of NF-κB, activator protein-1 (AP-1), and interferon regulatory factor (IRF) TF motifs underlying these state transitions. This coordinated TF activity may integrate persistent *α*-syn stimulation to drive progressive microglial activation and transition toward DAM. Consistent with this, microglial deletion of NF-κB p65 (RelA) or IRF4 impairs microglial progression toward DAM, with RelA governing early microglial activation and IRF4 regulating subsequent DAM differentiation. These TF motifs were also enriched in DAM in human PD, supporting conservation of this regulatory program across species. Together, our findings establish RelA and IRF4 as stage-specific regulators within the coordinated TF network that drives microglial state transitions toward DAM in PD.

## Results

### α-syn induces progressive microglial differentiation toward a functionally specialized DAM state

DAM have been implicated in multiple neurodegenerative disorders, including Alzheimer’s disease^14^ and PD^10,11^. Our prior work, using single-cell transcriptomics, identified an expansion of DAM in an *α*-syn overexpression mouse model of PD. However, it remains unclear whether this expansion is transient, or whether DAM represent a stable, functionally active microglial population that accumulates during disease progression. To address this, we utilized an adeno-associated virus (AAV)-mediated mouse model to drive neuronal *α*-syn overexpression (hereafter referred to as AAV2-SYN or AAV2-GFP as control) in the SNpc of *Spp1*-IRES-tdTomato reporter mice. We then assessed microglial states during peak and chronic neuroinflammation at 1 month and 3 months post-AAV2 transduction, respectively (**Fig. 1a**). *Spp1* has been previously found to be associated with the late DAM state^11,12^. To label this state without disrupting endogenous *Spp1* expression, we used *Spp1*-IRES-tdTomato knock-in reporter mice, in which tdTomato is inserted after the stop codon of *Spp1*. Based on the expression of the tdTomato reporter, we then categorized microglia into 4 major populations using established markers via flow cytometry: quiescent microglia (MHCII^-^), activated microglia (MHCII⁺ CD11c⁻ Spp1⁻), early DAM (MHCII⁺ CD11c⁺ Spp1⁻), and late DAM (MHCII⁺ CD11c⁺ Spp1⁺) (**Fig. 1b, Supplementary Fig. 1a**). Following *α*-syn overexpression, the proportion of microglia in activated and DAM states significantly increased in 3-month AAV2-SYN mice compared with 1-month AAV2-SYN mice, accompanied by a reduction in the quiescent population (**Fig. 1c**). At the protein level, changes in microglial states between AAV2-SYN and age-matched AAV2-GFP controls were significant at 3 months but not at 1 month. No proportional changes were observed between 1- and 3-month AAV2-GFP controls, indicating that these progressive microglial state transitions were driven by *α*-syn pathology. Consistent with their association with *α*-syn pathology, Clec7a^+^ Spp1^+^ DAM localized near *α*-syn-transduced neurons (**Supplementary Fig. 1c**).

**Fig. 1.**
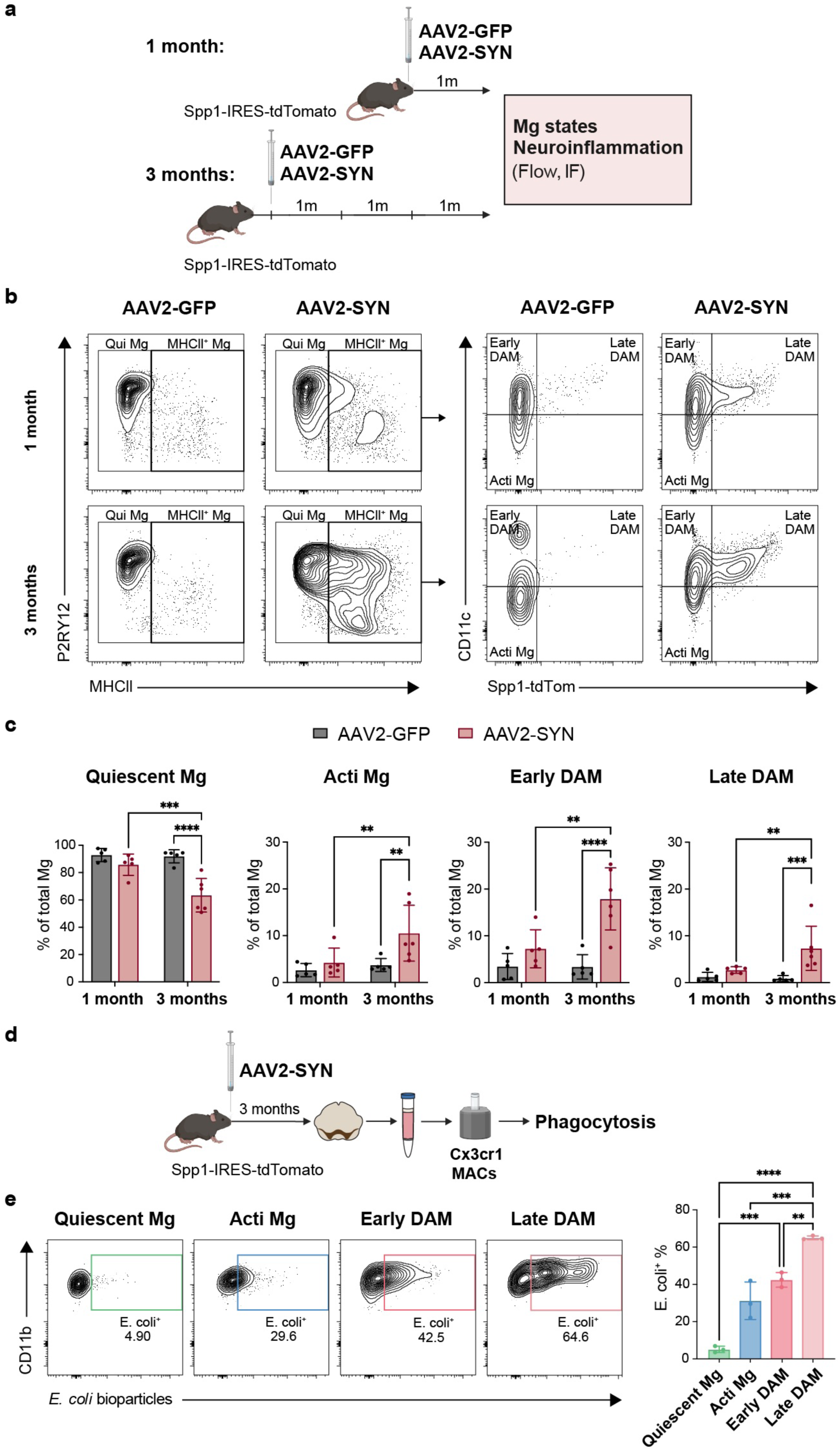
α-syn overexpression progressively drives microglia toward a functionally distinct DAM state. **a** Experimental schematic designed to assess microglial states and neuroinflammation in *Spp1*-IRES-tdTomato mice. 8-12-week-old mice were bilaterally injected with AAV2-GFP or AAV2-SYN into the SNpc. At 1 and 3 months post-AAV2 transduction, microglial states and peripheral immune cell infiltration were analyzed using flow cytometry. **b** Representative flow plots showing distinct microglial states. Microglia (Mg) were gated on live CD45^+^ CD11b^+^ Ly6C^-^ CD38^-^ P2RY12^+^ and further categorized as quiescent Mg (MHCII^-^), activated Mg (MHCII^+^), early DAM (MHCII^+^ CD11c^+^ Spp1^-^), and late DAM (MHCII^+^ CD11c^+^ Spp1^+^). **C** Quantification of the frequency of distinct microglial states out of total Mg at indicated timepoints. *n* = 5-6 per group. Two-way ANOVA, with Fisher’s LSD test. Mean ± SD are shown. **p ≤ 0.01, ***p ≤ 0.001, ****p ≤ 0.0001. **d** Experimental schematic of *ex vivo* phagocytosis assay using *E. coli* bioparticles in CX3CR1^+^ cells from *Spp1*-IRES-tdTomato mice at 3 months post-AAV2-SYN transduction. **e** Representative flow plots (left) and quantification (right) showing the frequency of *E. coli*^+^ cells across microglial states. *n* = 3 per group. One-way ANOVA, with Tukey’s test. Mean ± SD are shown. **p ≤ 0.01, ***p ≤ 0.001, ****p ≤ 0.0001. Panel **a** and **d** created in BioRender. Yang, V. (2026) https://BioRender.com/80l5wuz

We next sought to determine whether these microglial state transitions were associated with functional changes. We assessed key microglial functions, including phagocytosis and cytokine/chemokine production, across microglial states at 3 months post-AAV2-SYN transduction. Using an *ex vivo* phagocytosis assay, we found that the frequency of *E. coli*⁺ cells increased progressively across microglial states, with the highest phagocytic activity observed in late DAM (**Fig. 1d, e**). Similarly, *ex vivo* LPS and IFN-*γ* stimulation revealed that both early and late DAM exhibited significantly higher frequencies of TNF*α*, IL-1β, and CCL5-expressing cells than activated and quiescent microglia (**Supplementary Fig. 1d-g**). Together, these findings suggest that microglial state transitions toward DAM are accompanied by enhanced effector functions, identifying DAM as a functionally specialized microglial state.

Given their functional specialization, we next sought to determine the mechanisms underlying *α*-syn-induced microglial state transitions toward DAM. Because *α*-syn overexpression induces both microglial activation and peripheral immune responses, we first examined the contribution of infiltrating peripheral immune cells. We observed a significant decrease in monocyte/macrophage infiltration between 1- and 3-month AAV2-SYN mice (**Supplementary Fig. 1b**). While lymphocyte infiltration remained significantly elevated relative to AAV2-GFP controls, no further increase was observed between 1- and 3-month AAV2-SYN mice, suggesting that DAM accumulation is not accompanied by progressive peripheral immune cell infiltration. To further define the requirements for DAM formation, we examined Clec7a^+^ DAM in CD4 knockout and MHCII knockout mice at 3 month post-AAV2-SYN transduction, as CD4 T cells represent the majority of infiltrated lymphocytes in our model. While loss of CD4 T cells had no effect on DAM, knockout of MHCII greatly reduced DAM (**Supplementary Fig. 1h, i**), suggesting that microglial transitions toward DAM require MHCII-dependent microglial activation but occurs independently of CD4 T cell-mediated responses. Taken together, these findings demonstrate that *α*-syn overexpression drives progressive MHCII-dependent microglial transitions toward a functionally specialized DAM state characterized by enhanced phagocytosis and pro-inflammatory cytokine/chemokine production.

### A staged NF-κB–AP-1–IRF program underlies *α*-syn-induced chromatin remodeling and microglial reprogramming toward DAM

Given that MHCII deficiency suppresses DAM formation, we hypothesized that *α*-syn-induced microglial state transitions toward DAM are governed by microglia-intrinsic transcriptional and epigenetic programs. To identify these programs, we performed single-cell assay for transposase-accessible chromatin sequencing (scATAC-seq) on CX3CR1^+^ cells isolated at 1 month post-AAV2-GFP or AAV2-SYN transduction, a timepoint when DAM first emerge transcriptionally^11^ (**Fig. 2a**). Using Seurat, we integrated these datasets to specifically assess how *α*-syn overexpression changes chromatin accessibility in CNS-resident macrophages. Unbiased clustering identified 12 distinct clusters. As expected, border-associated macrophages (BAMs, cluster 7; *Pf4*, *Ccr2*, and *Mrc1*) were separated from microglial clusters (**Fig. 2b, c**). Based on differential chromatin accessibility profiles, microglial subclusters were categorized into distinct activation states, including quiescent microglia (clusters 3, 6, and 8; *Cx3cr1*, *Mafb*, and *Crybb1*), activated microglia (clusters 1, 2, and 9; *Atf3*, *Egr2*, *RelA* and *Nfkbia*), and DAM (clusters 4, 10, and 11; *Spp1*, *Clec7a*, and *Gpnmb*) (**Fig. 2c**). In addition, clusters 0 and 5 displayed increased chromatin accessibility associated with chromatin regulatory genes (*Phf19* and *Usp3)*, suggesting that they represent an intermediate chromatin-primed microglial state (hereafter referred to as primed microglia). Notably, clusters 11, 4, and 10 exhibited increased chromatin accessibility at distal regulatory regions associated with the *Spp1* and *Gpnmb* loci, consistent with activation of putative enhancers during *α*-syn-induced microglial state transitions (**Fig. 2d, Supplementary Fig. 2b**). To determine the trajectory of *α*-syn-induced microglial state transitions at the chromatin level, we performed pseudotime analysis with Monocle3 and identified a trajectory from quiescent and primed microglia through activated microglia and ultimately toward DAM, with DAM clusters 11, 4, and 10 likely representing progressive early, intermediate, and late DAM states, respectively (**Fig. 2e**). Although all clusters were present in both AAV2-GFP and AAV2-SYN experimental conditions, *α*-syn overexpression induced an expansion of clusters 4, 5, 10 and 11, with the most prominent expansion observed within DAM clusters (**Fig. 2f, g, Supplementary Fig. 2a**). Together, these findings suggest that *α*-syn overexpression establishes a more permissive chromatin landscape underlying microglial transitions toward DAM.

**Fig. 2.**
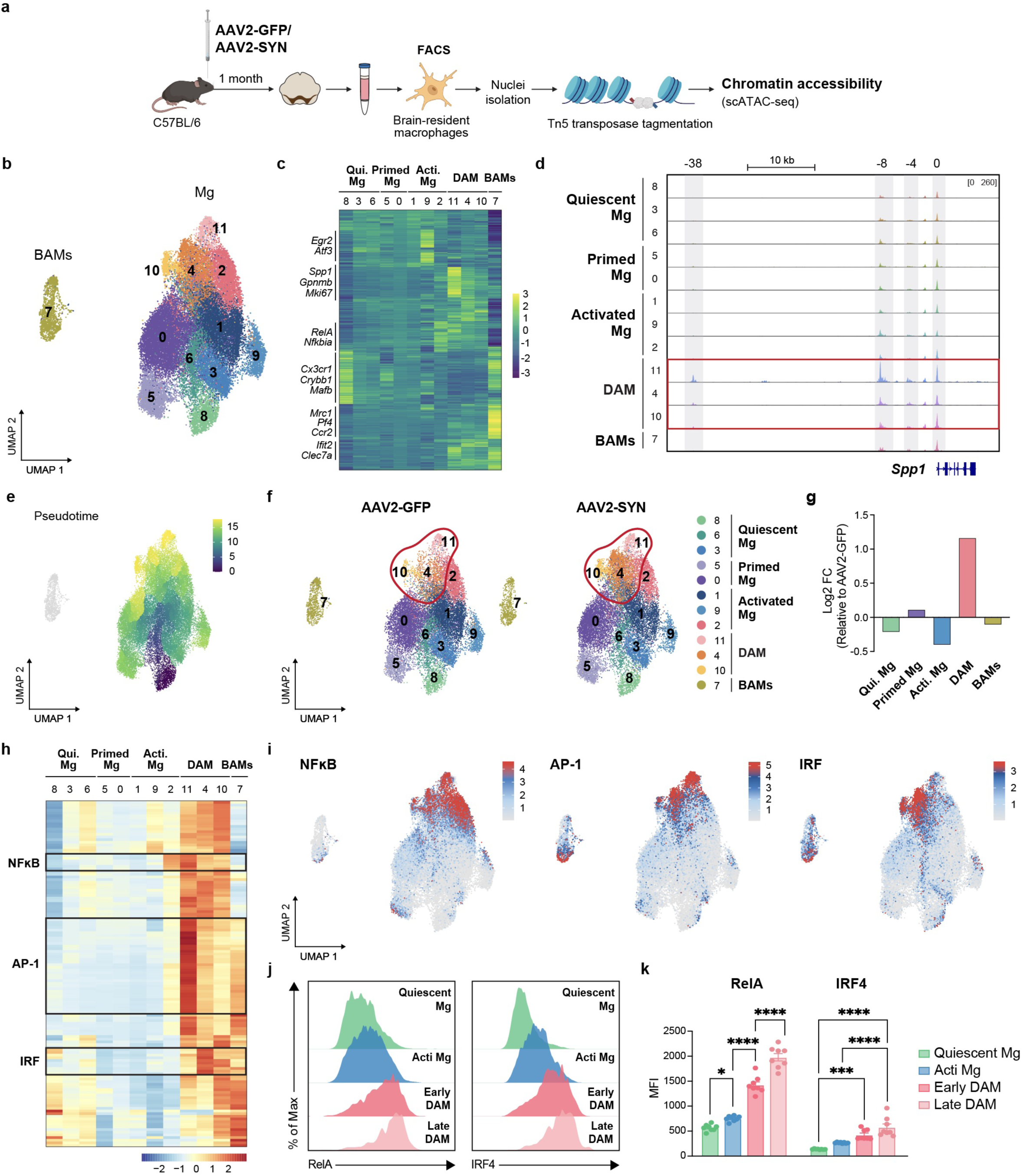
Increased chromatin accessibility at DAM-associated loci reveals a sequential NF-κB-AP-1-IRF axis driven by α-syn overexpression. **a** Experimental schematic of nuclei isolation from brain-resident macrophages for single-cell ATAC sequencing. 8-12-week-old C57BL/6 mice were bilaterally injected with AAV2-GFP or AAV2-SYN into the SNpc. At 1 month post-AAV2 transduction, live CD45^+^ CD11b^+^ CX3CR1^+^ Ly6C^-^ Ly6G^-^ NK1.1^-^ brain-resident macrophages were sorted for nuclei isolation. **b** Integrated UMAP plot showing 12 unique clusters. *n* = 2 per group, with 6 ventral midbrains pooled per *n*. **c** Heatmap showing log2 fold change in chromatin accessibility of each cluster in AAV2-SYN, compared to AAV2-GFP. Representative genes associated with increased chromatin accessibility are labeled. Cluster identities: 8, 3, and 6, Quiescent Mg; 5 and 0, Primed Mg; 1, 9, and 2, Activated Mg; 11, 4, and 10, DAM; 7, BAMs. **d** scATAC-seq peaks showing chromatin accessibility at the *Spp1* locus. Gray shaded boxes highlight increased accessibility at 0, -4, -8 and -38 kb before *Spp1* promoter in DAM clusters (11, 4, and 10). **e** Monocle pseudotime analysis overlaid on the integrated UMAP plot. **f** UMAP plots showing clusters from AAV2-GFP and AAV2-SYN. Red circles highlight changes in chromatin accessibility within DAM clusters (11, 4, and 10). **g** Bar graph showing log2 fold change of cluster groups in AAV2-SYN, compared to AAV2-GFP. **h** Heatmap showing the transcription factor (TF) motif enrichment score of each cluster in AAV2-SYN, compared to AAV2-GFP. Black boxes highlight motif enrichment from NF-κB, AP-1, and IRF families. **i** Integrated UMAP plots colored by enrichment of NF-κB, AP-1, and IRF families. NF-κB RelA and IRF4 protein expression in microglia were measured from *Spp1*-IRES-tdTomato mice at 1 month post-AAV2-SYN transduction by flow cytometry. **j** Representative histograms (**j**) and quantification (**k**) showing NF-κB RelA and IRF4 expression across microglial states, including quiescent Mg, activated Mg, early DAM, and late DAM. *n* = 8 per group. Two-way ANOVA, with Tukey’s test. Mean ± SEM are shown. *p ≤ 0.05, ***p ≤ 0.001, ****p ≤ 0.0001. Panel **a** created in BioRender. Yang, V. (2026) https://BioRender.com/hb3u0rw

Given that chromatin accessibility provides a regulatory landscape for transcription factor (TF) binding and gene regulation, we next sought to determine the TF regulatory programs underlying *α*-syn-induced microglial state transitions. To identify potential TFs associated with DAM differentiation, we performed chromVAR analysis to infer TFs activity based on chromatin accessibility. We observed a stepwise, overlapping pattern of TF motif enrichment along the microglial trajectory (**Fig. 2h**). NF-κB motif enrichment first increased in activated microglia (cluster 2) and peaked in early DAM (cluster 11), whereas AP-1 motifs peaked in early DAM (cluster 11) and remained enriched in intermediate DAM (cluster 4). IRF motifs subsequently peaked in intermediate DAM (cluster 4) and remained enriched in late DAM (cluster 10) (**Fig. 2i**). Consistent with *α*-syn’s known role in NF-κB activation^20^, these observations suggest a hierarchical TF regulatory program in which NF-κB activation precedes AP-1 and subsequent IRF programs during microglial reprogramming toward DAM. To further validate these motif predictions at the protein level, we measured TF protein expression across microglial states at 1 month post-AAV2-SYN transduction in *Spp1*-IRES-tdTomato reporter mice. We focused specifically on NF-κB p65 (RelA) and IRF4 because of their established roles in canonical NF-κB signaling and macrophage polarization, respectively^24,25^. We found that RelA expression progressively increased from quiescent to activated microglia, early DAM, and ultimately late DAM, whereas IRF4 expression was significantly higher in early and late DAM (**Fig. 2j, k**). These expression patterns were consistent with the observed TF motif enrichment. Taken together, these findings support a hierarchically organized TF program underlying *α*-syn-induced microglial state transitions, in which NF-κB contributes to early microglial activation and IRF4 to DAM differentiation.

### Microglial RelA or IRF4 deficiency impairs *α*-syn-induced microglial state transitions toward DAM

To determine whether RelA or IRF4 is required for *α*-syn-induced microglial state transitions toward DAM, we conditionally deleted *RelA* or *Irf4* in microglia. Tamoxifen-inducible *TMEM119*^CreERT^^2/CreERT2^ mice were crossed with *RelA*^fl/fl^ or *Irf4*^fl/fl^ to generate *TMEM119*^CreERT2/+^ *RelA*^fl/fl^ (*RelA* cKO) or *TMEM119*^CreERT2/+^ *Irf4*^fl/fl^ (*Irf4* cKO) mice. *TMEM119*^CreERT2/+^ mice were used as controls (Cre-ctrl). Cre recombination efficiency following tamoxifen treatment was approximately 90% in microglia, as determined in tamoxifen-treated *TMEM119*^creERT2/+^ tdTomato^fl+/-^ fate-mapping mice (**Supplementary Fig. 3a-c**). We evaluated different microglial states in Cre-ctrl, *RelA* cKO, and *Irf4* cKO mice at 1 month post-AAV2-SYN transduction via flow cytometry (**Fig. 3a**). Due to the absence of the *Spp1*-IRES-tdTomato reporter, microglia were categorized into quiescent (MHCII⁻), activated (MHCII⁺ CD11c⁻), and DAM (MHCII⁺ CD11c⁺) populations using a gating strategy similar to Fig. 1b. Compared with Cre-ctrl, *RelA* cKO and *Irf4* cKO mice showed a significant reduction in activated microglia and DAM, accompanied by an increase in quiescent microglia (**Fig. 3b, c**). Notably, the reduction in DAM was more pronounced in *RelA* cKO than in *Irf4* cKO, suggesting a greater contribution of NF-κB signaling during DAM differentiation. To further determine the functional consequence of impaired microglial state transitions toward DAM, we performed *ex vivo* phagocytosis assays on CX3CR1^+^ sorted cells isolated from Cre-ctrl, *RelA* cKO, and *Irf4* cKO mice at 1 month post-AAV2-SYN transduction (**Fig. 3d**). As expected, the number of phagocytosing microglia significantly decreased in *RelA* cKO and *Irf4* cKO compared with Cre-ctrl (**Fig. 3e, f**). Taken together, these findings demonstrate that RelA or IRF4 are required for *α*-syn-induced microglial state transitions toward the functionally specialized DAM state, with RelA deficiency producing a more pronounced reduction in DAM accumulation.

**Fig. 3.**
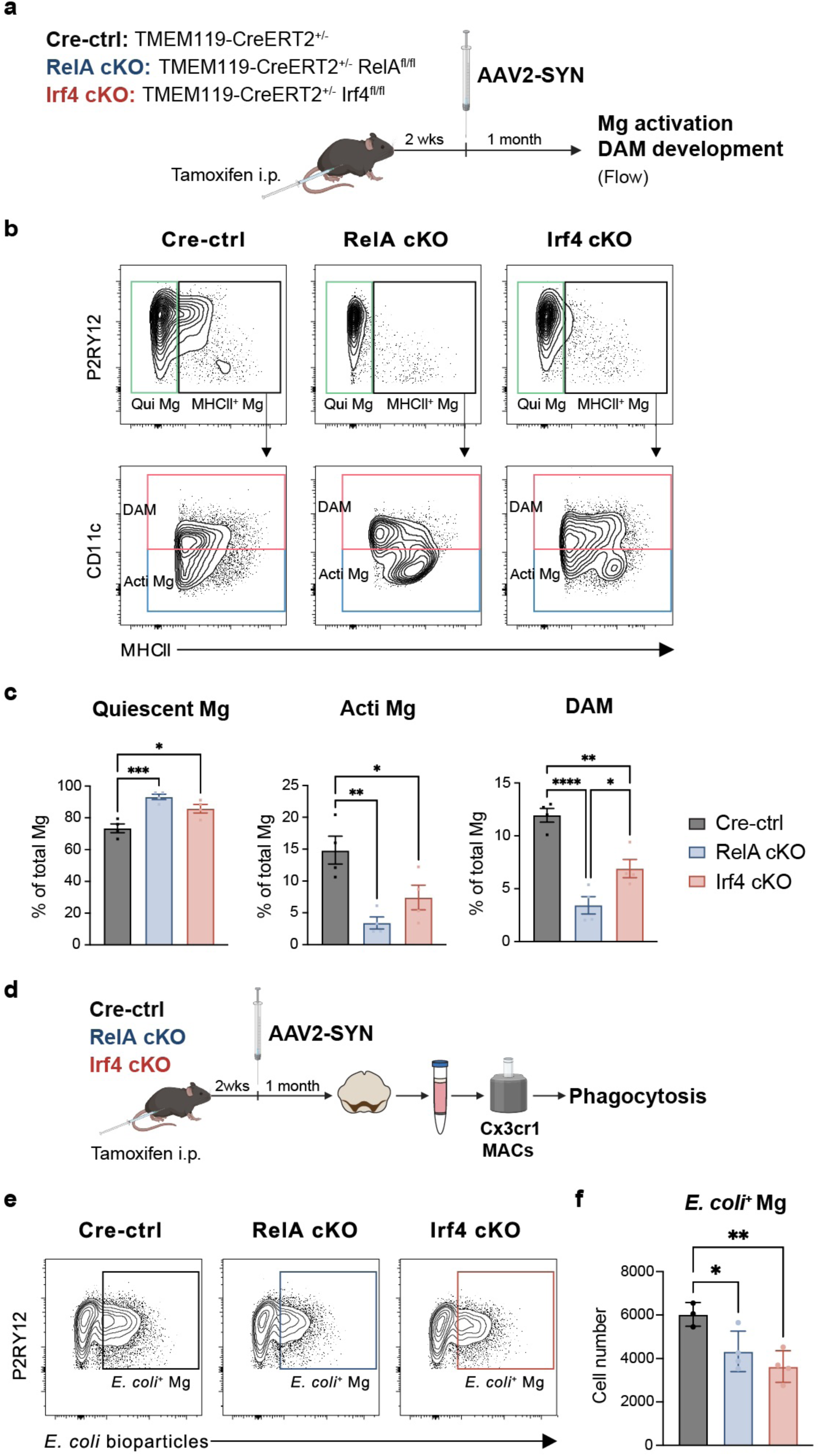
Conditional deletion of *RelA* or *Irf4* suppresses α-syn-induced DAM differentiation and phagocytosis. **a** Experimental schematic designed to assess distinct microglial states in Cre-ctrl (*TMEM119*-CreERT2^+/-^), *RelA* cKO (*TMEM119*-CreERT2^+/-^ *RelA*^fl/fl^), and *Irf4* cKO (*TMEM119*-CreERT2^+/-^ *Irf4*^fl/fl^) at 1 month post-AAV2-SYN transduction. Mice received tamoxifen, rested for 2 weeks, and were then bilaterally injected with AAV2-SYN. Microglia (Mg) were gated on live CD45^+^ CD11b^+^ Ly6C^-^ CD38^-^ P2RY12^+^ and further categorized as quiescent Mg (MHCII^-^), activated Mg (MHCII^+^ CD11c^-^), and DAM (MHCII^+^ CD11c^+^) using flow cytometry. **b, c** Representative flow plots (**b**) and quantification (**c**) showing frequency of quiescent Mg, activated Mg, and DAM out of total Mg. *n* = 4 per group, with 4 ventral midbrains pooled per *n*. One-way ANOVA, with Tukey’s test. Mean ± SEM are shown. *p ≤ 0.05, **p ≤ 0.01, ***p ≤ 0.001, ****p ≤ 0.0001. **d** Experimental schematic of *ex vivo* phagocytosis assay using *E. coli* bioparticles in CX3CR1^+^ cells isolated from Cre-ctrl, *RelA* cKO, and *Irf4* cKO at 1 month post-AAV2-SYN transduction. **e, f** Representative flow plots (**e**) and quantification (**f**) showing the absolute cell number of *E. coli*^+^ Mg in Cre-ctrl, *RelA* cKO, and *Irf4* cKO mice. *n* = 3-4 per group. One-way ANOVA, with Tukey’s test. Mean ± SEM are shown. *p ≤ 0.05, **p ≤ 0.01. Panel **a** and **d** created in BioRender. Yang, V. (2026) https://BioRender.com/hb3u0rw

### RelA and IRF4 shape stage-specific transcriptional programs during α-syn-induced microglial state transitions toward DAM

To determine how RelA and IRF4 shape the transcriptional programs underlying *α*-syn-induced microglial state transitions toward DAM, we performed single-cell RNA sequencing (scRNA-seq) on CX3CR1^+^ cells isolated from the ventral midbrains of Cre-ctrl, *RelA* cKO, and *Irf4* cKO mice at 1 month post-AAV2-SYN transduction (**Fig. 4a**). Using Seurat, we integrated Cre-ctrl, *RelA* cKO, and *Irf4* cKO datasets to specifically assess how RelA or IRF4 deficiency alters the transcriptional states of CNS-resident macrophages. Unbiased clustering identified 12 distinct clusters (**Fig. 4b)**. As expected, microglia and BAMs were separated into 2 distinct populations. Differential gene expression analysis identified clusters corresponding to BAMs (cluster 6, 8, 11; *Pf4* and *Mrc1*), quiescent microglia (clusters 1 and 2; *Sall1*, *Tmem119*, and *P2ry12*), activated microglia (clusters 0 and 3; *Atf3*, *Egr1* and *H2-Ab1*), proliferating microglia (clusters 7, 9, and 10; *Mki67* and *Top2a*), and DAM (clusters 4 and 5; *Itgax*, *Clec7a* and *Spp1*) (**Fig. 4c, d**). Consistent with our previous findings, label transfer from our published AAV2-GFP/AAV2-SYN dataset showed strong agreement in cluster identification^11^ (**Supplementary Fig. 4a**). To further examine microglial state transitions at the transcriptional level, we performed pseudotime analysis with Monocle3 and identified a microglial trajectory from quiescent to activated microglia, followed by branching toward proliferating microglia and DAM states (**Fig. 4e**). All clusters were present across the three experimental conditions. While both RelA and IRF4 deficiency shifted microglial states toward a more quiescent phenotype, they produced distinct alterations in state distribution (**Fig. 4f, Supplementary Fig. 4b**). In particular, RelA deficiency reduced activated microglia (cluster 3) and produced a more pronounced reduction in DAM clusters (clusters 5 and 4), whereas IRF4 deficiency primarily reduced DAM clusters (clusters 5 and 4). We also observed changes in other clusters, with IRF4 deficiency associated with a greater reduction in proliferating microglia and RelA deficiency associated with a greater reduction in BAMs. Together, these findings suggest distinct roles for RelA and IRF4 during *α*-syn-induced microglial state transitions toward DAM, with RelA acting at early activation states and IRF4 contributing more specifically to DAM differentiation.

**Fig. 4.**
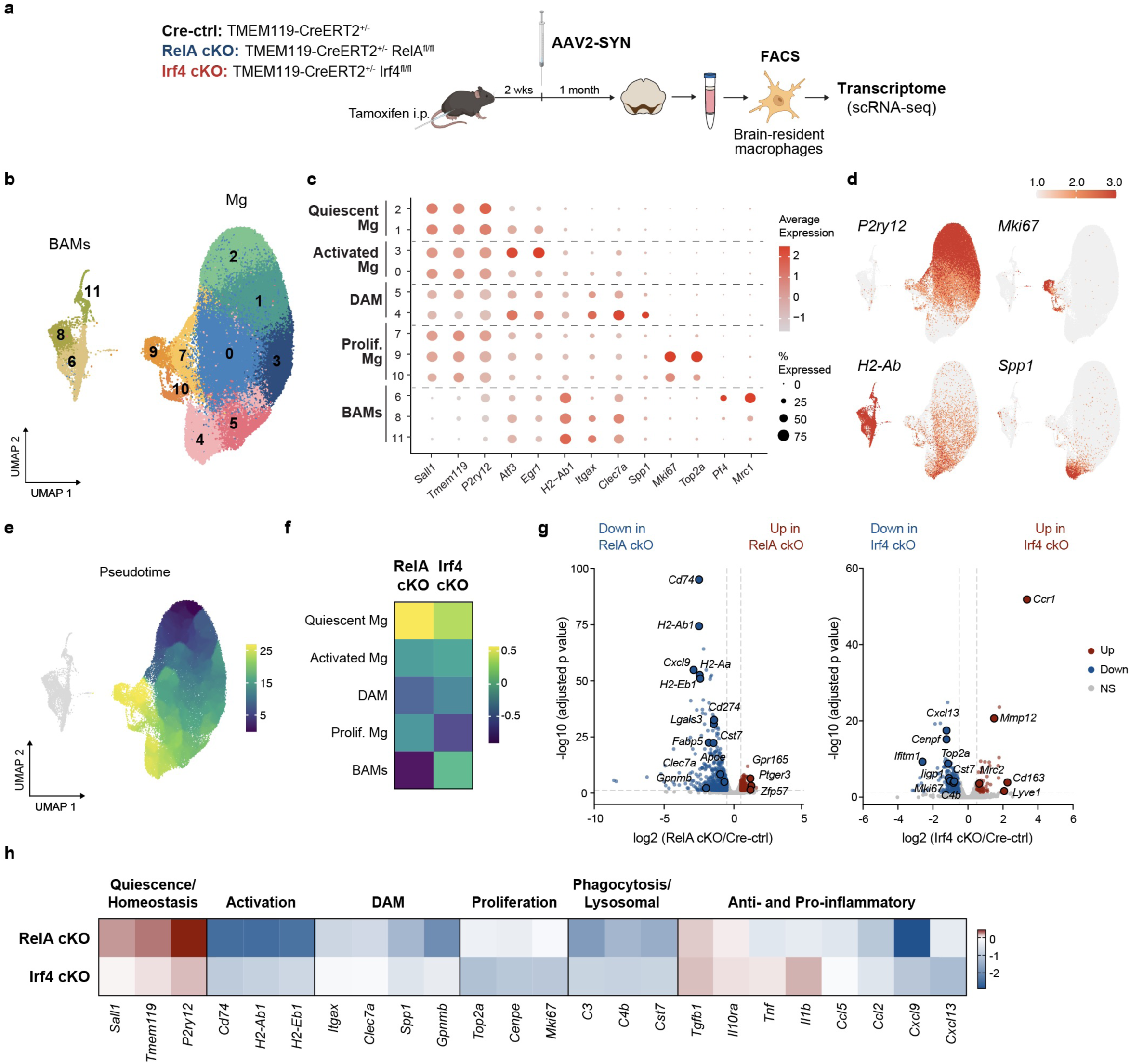
Distinct effects of RelA and IRF4 deficiency on α-syn-induced microglial activation, DAM differentiation, and proliferation. **a** Experimental schematic of brain-resident macrophages isolation for single-cell RNA sequencing. Cre-ctrl (*TMEM119*-CreERT2^+/-^), *RelA* cKO (*TMEM119*-CreERT2^+/-^ *RelA*^fl/fl^), and *Irf4* cKO (*TMEM119*-CreERT2^+/-^ *Irf4*^fl/fl^) mice were bilaterally injected with AAV2-SYN into the SNpc 2 weeks after tamoxifen administration. At 1 month post-AAV2 transduction, live CD45^+^ CD11b^+^ CX3CR1^+^ Ly6C^-^ Ly6G^-^ NK1.1^-^ brain-resident macrophages were sorted and processed for scRNA-seq. **b** Integrated UMAP plot showing 12 unique clusters. *n* = 2 per group, with 4 ventral midbrains pooled per *n*. **c** Dot plot showing cluster-defining genes for each cluster. Cluster identities: 2 and 1, Quiescent Mg; 3 and 0, Activated Mg; 5 and 4, DAM; 7, 9, and 10, Proliferating Mg; 6, 8, and 11, BAMs. **d** Integrated UMAP plot colored by representative cluster-defining genes (*P2ry12*, *Mki67*, *Spp1*, *H2-Ab1*). **e** Monocle pseudotime analysis overlaid on the integrated UMAP plot. **f** Heatmap showing log2 fold change of cluster groups in *RelA* cKO and *Irf4* cKO, compared to Cre-ctrl. **g** Volcano plot showing log2 fold change in gene expression in *RelA* cKO (left) and *Irf4* cKO (right), compared to Cre-ctrl. Representative genes that were significantly different were labeled. **h** Heatmap showing representative genes for quiescence/homeostasis, activation, DAM, proliferation, phagocytosis/lysosomal function, and inflammation in *RelA* cKO and *Irf4* cKO, compared to Cre-ctrl. Panel **a** created in BioRender. Yang, V. (2026) https://BioRender.com/hb3u0rw

To further define the transcriptional programs altered by RelA and IRF4 deficiency, we performed pseudobulk differential expression analysis. We found significant downregulation of genes associated with microglial activation, antigen presentation, and DAM-associated programs (*H2-Ab1*, *Cd74*, *Lgals3*, *Clec7a*) in *RelA* cKO compared with Cre-ctrl (**Fig. 4g**), consistent with the reduced activated microglia and DAM states in *RelA* cKO. Genes associated with interferon responses and proliferation (*Ifitm1*, *Top2a*, *Mki67*) were significantly decreased in *Irf4* cKO compared with Cre-ctrl (**Fig. 4g**). As DAM differentiation is accompanied by the acquisition of effector programs, we next examined genes involved in phagocytosis/lysosomal pathways and inflammatory signaling. Both RelA and IRF4 deficiency downregulated phagocytic and pro-inflammatory genes (*C3*, *Cst7*, *Ccl5*, *Cxcl9*) while increasing genes associated with immunoregulatory signaling (*Tgfb1* and *Il10ra*) (**Fig. 4h**), suggesting that RelA and IRF4 are critical regulators of DAM-associated functional programs. Taken together, these findings indicate that RelA and IRF4 coordinate a temporally organized TF program that drives *α*-syn-induced microglial state transitions toward a functionally specialized DAM state, with RelA shaping early microglial activation and IRF4 regulating subsequent DAM differentiation.

### The NF-κB–AP-1–IRF program is conserved during microglial progression toward DAM in human PD

Given that the NF-κB–AP-1–IRF program regulates microglial reprogramming toward DAM in our *α*-syn overexpression mouse model, we next asked whether this TF program is conserved during DAM differentiation in human PD. Using a previously published single-nucleus RNA-seq dataset from the substantia nigra (13 PD and 15 control samples)^26^, we subclustered human microglia and identified 4 major populations based on their marker gene expression: quiescent microglia (cluster 3), surveillant microglia (cluster 2), activated microglia (cluster 0), and DAM (cluster 1) (**Fig. 5a-c**). In human PD, the DAM cluster expressed canonical DAM-associated genes such as *SPP1*, *APOE*, *CLEC7A*, and *GPNMB* (**Fig. 5d**), consistent with the DAM-associated transcriptional profiles observed in the mouse model. To determine whether the transcriptional signature of the late DAM state identified in our mouse model is conserved in human PD, we calculated module scores for the mouse late DAM signature across human microglial clusters. The mouse late DAM signature progressively increased across microglial activation states and displayed the highest score in the human DAM cluster (**Fig. 5e**). Consistent with our finding that *α*-syn overexpression promotes microglial state transitions toward DAM, the mouse late DAM signature scores within the human activated microglia and DAM clusters were significantly elevated in PD tissue compared to controls (**Fig. 5f**). Together, these findings indicate that human PD exhibits increased DAM transcriptional signatures similar to those identified in our mouse model.

**Fig. 5.**
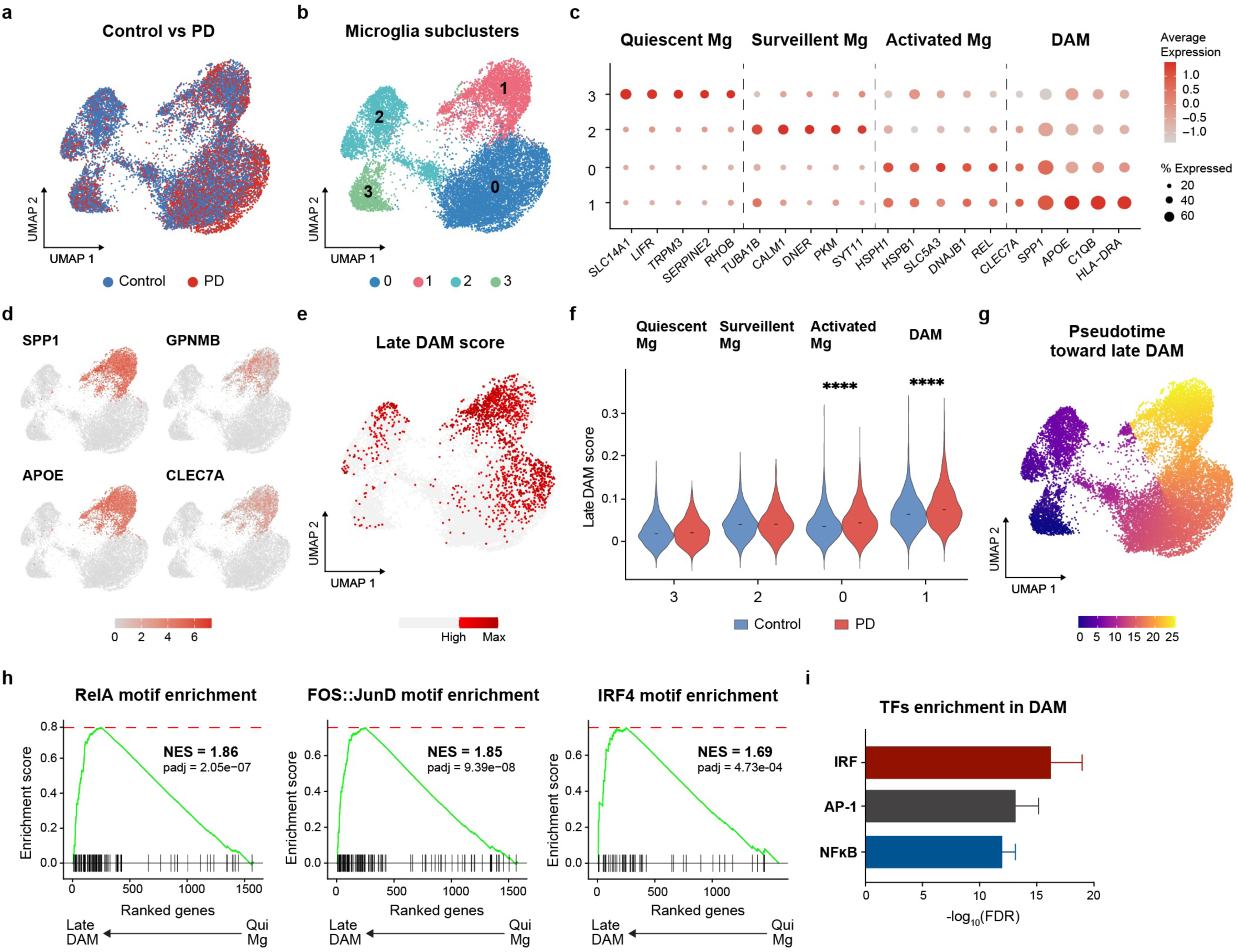
Human PD exhibits increased DAM and enriched NF-κB/AP-1/IRF signaling along the DAM trajectory. **a** UMAP plot showing nuclei from the substantia nigra of control (n = 15) and PD (n = 13). **b** Integrated UMAP plot of substantia nigra microglial nuclei identifying 4 unique clusters. **c** Dot plot showing cluster-defining genes for each microglial cluster. Cluster identities: 3, Quiescent microglia; 2, Surveillant microglia; 0, Activated microglia; 1, DAM. **d** Integrated UMAP plots colored for expression of DAM genes. **e** Integrated UMAP plot colored by mouse late DAM score, calculated using AddModuleScore. **f** Quantification of mouse late DAM score in human microglial clusters in control and PD. Two-tailed Wilcoxon test. Median is shown. ****p ≤ 0.0001. **g** Pseudotime analysis showing microglial trajectory toward the late DAM state overlaid on the integrated UMAP plot. **h** Pre-ranked gene set enrichment analysis of gene sets associated with the RelA, FOS::JunD (AP-1), and IRF4 motifs, using genes ranked by their expression change along pseudotime. Normalized enrichment scores (NES) and adjusted two-sided p-values are indicated. **i** Enrichr analysis showing enrichment of NFκB, AP-1, and IRF families in human DAM. -log10 (false discovery rate) is plotted. Mean ± SEM are shown.

To further characterize the transcriptional progression toward DAM in human PD, we performed pseudotime analysis with Slingshot. Because human DAM exhibited the highest mouse late DAM signature, we defined DAM as the terminal state, and identified a trajectory from quiescent to surveillant to activated microglia, and ultimately to DAM (**Fig. 5g**). To determine whether the NF-κB–AP-1–IRF program is conserved during this progression, we performed gene set enrichment analysis along the microglial trajectory, using RelA, Fos:JunD, and IRF4-associated gene programs derived from our mouse scATAC-seq analysis. Significant enrichment of RelA, Fos:JunD, and IRF4-associated gene programs was observed along the trajectory toward DAM, suggesting conservation of these TF-associated programs during human DAM differentiation (**Fig. 5h**). To independently assess TF enrichment in human DAM, we performed TF enrichment analysis using ChEA3. Consistent with our GSEA results, NF-κB, AP-1, and IRF family TFs were significantly enriched in human DAM (**Fig. 5i, Supplementary Table 1**). Collectively, these findings support conservation of the NF-κB–AP-1–IRF regulatory program during microglial progression toward DAM across the *α*-syn overexpression mouse model and human PD.

## Discussion

While DAM have emerged as a conserved microglial state across neurodegenerative disorders, the transcriptional and epigenetic mechanisms that drive their differentiation remain poorly understood. Here, we identified a temporally coordinated NF-κB–AP-1–IRF regulatory program that accompanies *α*-syn-induced microglial state transitions toward DAM, with NF-κB p65 (RelA) and IRF4 exerting distinct, stage-specific roles in early microglial activation and subsequent DAM differentiation. Disruption of this program impairs progression toward the DAM state, while enrichment of NF-κB, AP-1, and IRF motifs in DAM from human PD supports conservation of this regulatory program across species.

Within, we investigated the transcriptional regulatory networks and factors governing microglial state transitions toward DAM. Our integrative transcriptomic and chromatin accessibility analyses revealed a coordinated temporal sequence, overlapping enrichment of NF-κB, AP-1, and IRF motifs progressing from activated microglia to early DAM and ultimately to late DAM. This was accompanied by increased chromatin accessibility at distal regulatory regions of DAM-associated loci, suggesting that chronic *α*-syn stimulation promotes chromatin remodeling and *de novo* enhancer establishment during microglial state transitions toward DAM. Given that *α*-syn engages TLR2/4 and activates NF-κB^20^, and that NF-κB can cooperate with AP-1 to remodel regulatory landscapes during sustained TLR signaling^22,27,28^, early NF-κB activity may establish a regulatory environment that enables subsequent IRF-dependent DAM differentiation. Thus, chronic neurodegenerative stimuli such as *α*-syn may progressively build the regulatory landscape of the DAM state through sequential engagement of NF-κB–AP-1–IRF transcriptional regulators.

Despite identification across multiple neurodegenerative disorders, it remained unclear whether a shared or disease-specific transcriptional regulatory program governs DAM differentiation. The enrichment of NF-κB, AP-1, and IRF family TFs in DAM in both our *α*-syn mouse model and human PD supports conservation of this program across species. Moreover, studies in amyloid-β models of Alzheimer’s disease have identified DAM-associated TFs from the same families, including RelA^29,30^, raising the possibility that the NF-κB–AP-1–IRF axis represents a shared regulatory framework through which diverse neurodegenerative stimuli drive microglial state transitions. At the same time, other DAM-associated TFs, including MITF and BHLHE40/41 in AD and NR4A2 and Bach1 in PD, may provide disease- or context-specific inputs. Together, these observations suggest that diverse neurodegenerative stimuli may converge on a conserved regulatory program while engaging additional factors that shape microglial state in a disease- and context-dependent manner.

While our findings define the transcriptional and epigenetic programs that drive DAM differentiation, the functional consequences of this specialized state remain unclear, with evidence for both protective and detrimental effects depending on context^31–33^. In this study, DAM exhibited increased phagocytic activity and enhanced cytokine and chemokine production upon stimulation, consistent with a functionally primed state capable of mounting rapid responses to pathological cues. Such priming may be beneficial when microglia can effectively clear accumulating pathological material, however this may become maladaptive when persistent or excessive stimulation exceeds their capacity to resolve the underlying pathology. The frequent accumulation of DAM around *α*-syn and amyloid-β pathology is consistent with their engagement in the response to pathological material, although whether DAM-mediated activity is sufficient to promote effective clearance remains unclear. Conversely, heightened production of inflammatory mediators such as TNFα, IL-1β, and CCL5 could amplify local inflammation and recruit peripheral immune cells, potentially contributing to tissue injury. Thus, rather than being intrinsically protective or detrimental, DAM may represent a conserved adaptation to chronic stimulation whose functional consequences depend on the balance between pathological demand and the capacity of the microglial response to resolve it. Future studies defining the cellular and molecular signals that determine DAM function *in vivo* will be important for defining the role of DAM in PD and other neurodegenerative disorders.

The persistence of DAM following individual TF perturbations highlights the importance of this regulatory response. Although microglia-specific deletion of *RelA* or *Irf4* significantly impaired DAM progression at both the transcriptional and protein levels, neither perturbation completely abolished the DAM state. This residual differentiation may reflect functional compensation among related TFs, as members of the NF-κB and IRF families share overlapping DNA-binding properties and can regulate common target genes^34,35^. Such redundancy may provide regulatory robustness that allows microglia to maintain adaptive responses despite disruption of individual pathway components, potentially preserving DAM-associated functions during persistent neurodegenerative stress or cues. This redundancy also means that single-TF perturbations are unlikely to fully capture the contribution of the DAM program, particularly as microglial states continue to evolve during prolonged stimulation or over time. Finally, although DAM exhibited enhanced phagocytic activity and localized near *α*-syn pathology, the high and relatively acute *α*-syn burden produced by viral overexpression limits our ability to determine whether this response is sufficient to promote physiological *α*-syn clearance. Future studies using combinatorial TF perturbation and models that more gradually recapitulate pathological protein accumulation will be important for determining how the conserved DAM response contributes to maintaining microglial function and controlling pathology *in vivo*.

In summary, our findings identify a conserved NF-κB–AP-1–IRF regulatory network through which chronic *α*-syn stimulation progressively establishes the DAM state. RelA and IRF4 act at distinct stages of this process, linking early microglial activation to subsequent DAM differentiation and regulatory remodeling. The conservation of this architecture across human PD and other neurodegenerative contexts suggests that DAM represent a broadly conserved microglial adaptation to persistent innate immune stimulation. This adaptation may enhance the capacity of microglia to respond to pathological proteins, while its ultimate consequences depend on the intensity and persistence of the signals encountered after differentiation.

## Methods

### Mice

C57BL/6J, (#000664, The Jackson Laboratory), C57BL/6J-*Spp1*^em1Msasn^/J, referred to as *Spp1*-IRES-tdTomato (#033731, The Jackson Laboratory), C57BL/6-*Tmem119*^em1(cre/ERT2)Gfng^/J, referred to as *TMEM119*-CreERT2 (#031820, The Jackson Laboratory), B6.Cg-*Gt(ROSA)26*Sor^tm9(CAG-tdTomato)Hze^/J, referred to as tdTomato (#007909, The Jackson Laboratory), B6.129S1-*Rela*^tm1Ukl^/J, referred to as *RelA*^fl/fl^ (#024342, The Jackson Laboratory), and B6.129S1-*Irf4*^tm1Rdf^/J, referred to as *Irf4*^fl/fl^ (#009380, The Jackson Laboratory) were used for this study. Mice were bred and maintained on a congenic background. Both males and females were utilized in all studies. All research conducted on animals was approved by the Institutional Animal Care and Use Committee at the University of Alabama at Birmingham.

### AAV2-GFP and AAV2-SYN virus

Green fluorescent protein or human full-length *α*-syn neuronal overexpression was mediated by recombinant adeno-associated virus, serotype 2 (AAV2). Manufacturing, construction and purification of the AAV2 vectors, AAV2/1-CBA-IRES2eGFP-WPRE (CIGW, AAV2-GFP) and AAV2/1-CBA-haSYN-IRES2eGFP-WPRE (CISGW, AAV2-SYN), were performed by the University of Iowa Viral Vector Core as previously described^36,37^.

### Stereotaxic surgery

Surgery was performed as previously described^37^. Briefly, mice were deeply anesthetized with inhaled 3% isoflurane and immobilized in a stereotaxic frame. The mice were injected stereotaxically with 2 μL of AAV2-SYN (9.21 × 10^11^ to 1.5 × 10^12^ viral genomes/mL) or AAV2-GFP (at equivalent titers to AA2-SYN) diluted in saline into the substantia nigra pars compacta (SNpc) at a rate of 0.5 µL/min with Hamilton syringe. After each injection, the needle was left in the SNpc for at least 2 minutes, and then slowly withdrawn. The stereotaxic coordinates for the SNpc were anterior-posterior −3.2 mm from bregma, mediolateral ±1.2 mm from midline, and dorsoventral −4.6 mm from dura. Mice for immunohistochemistry and unbiased stereology were unilaterally injected, while mice for flow cytometry and single-cell sequencing were bilaterally injected. For All surgical procedures and postoperative care protocols were followed and approved by the Institutional Animal Care and Use Committee at the University of Alabama at Birmingham.

### Tamoxifen treatment

For specific *RelA* or *Irf4* deletion in microglia, tamoxifen-inducible *TMEM119*-CreERT2 were crossed with *RelA*^fl/fl^ or *Irf4*^fl/fl^ to generate *TMEM119*-CreERT2^+/-^ *RelA*^fl/fl^ or *TMEM119*-CreERT2^+/-^ *Irf4*^fl/fl^. For genetic control, *TMEM119*-CreERT2 were crossed with C57BL/6J to generate *TMEM119*-CreERT2^+/-^. To bypass potential developmental deficits of gene deficiency, 6-8 weeks old of *TMEM119*-CreERT2-positive mice were injected with 3 mg tamoxifen dissolved in 100 μL corn oil daily intraperitoneally for 5 consecutive days. Mice rested for 2 weeks after tamoxifen treatment before AAV2 injection to allow stable recombination and tamoxifen clearance. At this time point, fate-mapping *TMEM119*-CreERT2^+/-^ tdTomato mice were used to determine recombination efficiency for microglia (∼90%), BAMs (∼26%), and monocytes (∼0%).

### Tissue preparation and processing

1, 3 or 6 months post-viral transduction, mice were deeply anaesthetized and transcardially perfused with a 0.1% heparinized PBS, followed by 4% paraformaldehyde (PFA) in PBS. Brains were collected and post-fixed in 4%PFA for 3 hours and transferred to 30% sucrose in PBS at 4°C for cryoprotective for over 48 hours. Brains were frozen on dry ice and sectioned coronally at a 40-μm thickness using a sliding microtome. Sectioned brains were serially collected and stored in 50% glycerol in PBS at -20°C.

### Immunohistochemistry

For fluorescent staining, free-floating sections were washed with 1X Tris-buffered saline (TBS), blocked with 5% serum from the secondary antibody host species in TBS for 1 hour, and incubated with primary antibodies in 1% serum from the same species in TBS containing 0.1% Triton X-100 (TBST) overnight at 4°C. The following primary antibodies were used: anti-α-synuclein (phospho-Serine129, clone EP1536Y, Abcam, ab51253, 1:5000), anti-IBA1 (polyclonal, Wako, NC1718288, 1:800), anti-dectin1 (clone R1-8g7, InvivoGen, mabg-mdect, 1:40), anti-MHCII (clone M5/114.15.2, eBiosciences, 50-112-9455, 1:500). Sections were washed and incubated with appropriate Alexa-fluor conjugated secondary antibodies diluted in 1% serum in TBST for 2 hours. The following cross-absorbed fluorescent conjugated secondaries were used: Alexa Fluor 555 goat anti-rat (Invitrogen, A21434, 1:1000), Alexa Fluor 647 goat anti-rabbit (Invitrogen, A21244, 1:1000), Alexa Fluor 647 goat anti-rat (Invitrogen, A21247, 1:1000). Sections were then washed, mounted on coated glass slides, and coversliped with HardSet antifade mounting medium (Vector Laboratories, H1500-10) to preserve fluorescent signals. Florescent microscopic images were acquired using a Nikon AX Confocal microscope. Images were then exported and processed using Adobe Photoshop and Adobe Illustrator. For quantification, 2-3 sections per brain encompassing the SNpc were imaged and quantified using Fiji/ImageJ 1.54p and representative images were selected.

For diaminobenzidine (DAB) staining, free-floating sections were washed with TBS, quenched of endogenous peroxidases with 3% hydrogen peroxide in TBS, blocked with 5% serum from the secondary antibody host species in TBS for 1 hour, and incubated with primary antibodies in 1% serum from the same species in TBST overnight at 4°C. The following primary antibodies were used: anti-α-synuclein (phospho-Serine129, clone EP1536Y, Abcam, ab51253, 1:2000), anti-IBA1 (polyclonal, Wako, NC1718288, 1:500), anti-dectin1 (clone R1-8g7, InvivoGen, mabg-mdect, 1:40), anti-MHCII (clone M5/114.15.2, eBiosciences, 50-112-9455, 1:500), anti-tyrosine hydroxylase (TH, polyclonal, Sigma-Aldrich, AB152, 1:2000). Sections were washed and incubated with appropriate biotinylated secondary antibodies diluted in TBST for 2 hours. The following biotinylated secondaries were used: Biotinylated goat-anti rabbit (Invitrogen, 65-6140, 1:1000), Biotinylated goat-anti rat (Vector Laboratories, BA-9401, 1:1000). Sections were then washed, incubated with VECTASTAIN ABC Reagent (Vector Labs, PK7100) for 30 minutes and developed with DAB Substrate Kit (Vector Labs, SK4100) according to the manufacturer’s instructions. Sections were washed, mounted on coated glass slides, dehydrated and coverslipped with Permount mounting medium (Electron Microscopy Sciences, 17986-01). Brightfield images were acquired and exported using a Zeiss Imager M2 brightfield microscope (MBF Biosciences). For quantification, 3 sections per brain encompassing the SNpc were imaged and quantified using Fiji/ImageJ 1.54p and representative images were selected.

### Unbiased stereology

For quantification of TH neurons in the SNpc, unbiased stereology was performed as previously described^38,39^. In brief, TH-DAB stained SNpc slides were coded and analyzed by a blinded reviewer using a Zeiss Axio Imager M2 microscope with StereoInvestigator Software (MBF Biosciences). 5-6 sections per brain encompassing the SNpc were were quantified using the optical fractionator method. TH-positive neurons within the contours of the SNpc were counted on a 100 μm × 100 μm grid with a 50 μm × 50 μm counting frame and an optical dissector height of 22 μm. Weighted section thickness was used to correct variations for tissue thickness. Representative brightfield images depicting TH neurons in the SNpc were acquired with Zeiss Axio Imager M2 microscope.

### Mononuclear cell isolation

1 or 3 months post transduction, mice were deeply anaesthetized and transcardially perfused with 1X DPBS without calcium and magnesium. Mononuclear cells within the ventral midbrain were isolated as previously described. Briefly, 3-mm section of midbrain was dissected, dissociated and digested with 1 mg/mL Collagenase IV (Sigma, C5138) diluted in RPMI 1640 (Gibco, 11875093) at 37°C. Digested cells were triturated and passed through 70-μm strainers to obtain single-cell suspension. Mononuclear cells were isolated using a 30/70% Percoll gradient (Cytiva, 17089101) and the resulting interphase layer was collected.

### Flow cytometry

Isolated mononuclear cells were blocked with rat anti-CD16/32 (553141, BD Biosciences, 1:100). Cells were washed and labeled with the following antibodies for 30 min at 4°C: anti-CD45 BUV661 (clone 30-F11, BD Biosciences, 612975, 1:400), anti-CD45 BUV737 (clone I3/2.3, BD Biosciences, 752414, 1:200), anti-CD45 BUV805 (clone 30-F11, BD Biosciences, 568336, 1:200), anti-Ly6C BUV805 (clone HK1.4, BD Biosciences, 755202, 1:200), anti-CD11b e450 (clone M1/70, eBioscience, 48-0112-82, 1:200), anti-CD11c BV421 (clone N418, Biolegend, 117329, 1:50), anti-CD11b BV605 (clone M1/70, Biolegend, 101257, 1:100), anti-CD11c BV605 (clone N418, Biolegend, 117333, 1:100), anti-PD-L1 BV711 (clone 10F.9G2, Biolegend, 124319, 1:50), anti-MHCII BV711 (clone M5/114.15.2, Biolegend, 107643, 1:250), anti-MHCII BV785 (clone M5/114.15.2, Biolegend, 107645, 1:250), anti-CD11c BV785 (clone N418, Biolegend, 117336, 1:50), anti-Ly6C BV785 (clone HK1.4, Biolegend, 128041, 1:200), anti-TCRβ BV785 (clone H57-597, Biolegend, 109249, 1:200), anti-P2RY12 AF488 (clone S16007D, Biolegend, 848016, 1:200), anti-CD38 PeCy7 (clone 90, Biolegend, 102718, 1:200), anti-Ly6C APC (clone HK1.4, Biolegend, 128015, 1:200), anti-CD38 APC (clone 90, Biolegend, 102712, 1:200). A fixable viability dye was also used according to manufacturer’s instructions (eBioscience, 65-0865-14, 1:800).

For intracellular TFs staining, isolated mononuclear cells were Fc blocked, surface stained, processed using the Foxp3/Transcription Factor Staining Kit (eBioscience, 00-5523-00), and labeled with the following antibodies for 45 min at room temperature: anti-Sall1 e450 (clone NRNSTNX, eBioscience, 48-9729-82, 1:100), anti-NF-κB p65 Pacific Blue (clone D14E12, Cell Signaling Technology, 18145S, 1:100) and anti-IRF4 PeCy7 (clone 3E4, eBioscience, 25-9858-82, 1:100).

For intracellular cytokine/chemokine staining, isolated mononuclear cells were first stimulated with LPS (Invivogen, tlrl-3pelps, 1 μg/mL) and IFN-*γ* (R&D, 485-MI, 10 ng/mL) in the presence of GolgiStop (BD Biosciences, 554724, 1:1000) and GolgiPlug (BD Biosciences, 555029, 1:1000) for 4 hours at 37°C/5% CO_2_. Cells were then Fc blocked, surface stained, processed using the Foxp3/Transcription Factor Staining Kit, and labeled with the following antibodies for 45 min at room temperature: anti-TNF FITC (clone MP6-XT22, eBioscience, 11-7321-81, 1:50), anti-CCL5 PeCy7 (clone 2E9/CCL5, Biolegend, 149105, 1:75), anti-IL1β APC (clone NJTEN3, eBioscience, 17-7114-80, 1:100). Samples were run on a BD Symphony (BD Biosciences) and analyzed using FlowJo™ v10.10.0 Software (BD Life Sciences).

### *ex vivo* phagocytosis assay

CX3CR1^+^ cells were enriched from isolated mononuclear cells using MojoSort™ Mouse CX3CR1 Selection Kit (Biolegend, 480056) according to according to manufacturer’s instructions. Enriched CX3CR1^+^ cells were counted using a hemocytometer and incubated with pHrodo™ Deep Red E. coli BioParticles™ (Invitrogen™, P35360) at a particle-to-cell ratio of 10:1 at 37°C for 15 minutes. Cells were immediately placed on ice to terminate phagocytosis and proceeded with Fc block and surface antibody staining. Phagocytic activity was analyzed using flow cytometry.

### Single-cell RNA sequencing (scRNA-seq)

Mononuclear cells were isolated for sequencing as described above with 4 ventral midbrains pooled per sample, and sorted for CD45^+^ CD11b^+^ CX3CR1^+^ Ly6C^-^ Ly6G^-^ and NK1.1^-^ cells on a BD FACsAria. Sorted cells were loaded onto the 10X Chromium platform (10x Genomics), and libraries were constructed using the Next GEM Single Cell 3’v4 Kit according to the manufacturer’s instructions. Two biological replicates for each group were processed separately. Samples were sequenced on an Illumina NovaSeq at an average depth of 20,000 reads per cell. Sequencing files were processed, mapped to mm10, and count matrices were extracted using the Cell Ranger Single Cell Software (v 9.0.0).

### scRNA-seq analysis

Analyses were performed in R using Seurat (v3.1). Data was pre-processed by removing genes expressed in fewer than 3 cells and excluding cells that were outliers for the number of RNA molecules or more than 12% mitochondrial genes. The datasets were merged together and integrated following the Seurat standard integration method using the 2000 most variable genes. Following normalization, UMAP dimensional reduction was performed using the first 20 principal components. Clustering was performed following identification of nearest neighbors, using the first 20 dimensions and a resolution of 0.7. Clusters predominantly composed of either cells with low RNA content, doublet cells, or non-macrophage lineage cells were removed and the datasets were re-clustered following dimensional reduction. Marker genes for each cluster were determined using the FindAllMarkers function of Seurat with a minimum Log2 fold change threshold of +/−0.25 with the Wilcoxon ranked-sum test.

### Pseudotime analysis

Filtered and merged datasets were imported into Monocle3 by generating a cell dataset from the raw counts slot of the Seurat object on the 2000 most variable genes. Normalization and PCA were done with the preprocess_cds command from Monocle3 using the first 50 dimensions, and batch correction was applied using the align_cds command, which utilizes the Batchelor tool (Haghverdi et al.39). UMAP dimensional reduction was performed using the reduce_dimension command. Cells were clustered with cluster_cells using “Louvain” with the k set to 20. The trajectory graph was learned on the Monocle-derived clusters by calling learn_graph. Cells on the UMAP plot are colored by Seurat-derived clusters. Pseudotime was determined using the quiescent clusters as the starting point.

### Single-cell ATAC sequencing (scATAC-seq)

Mononuclear cells were isolated for sequencing as described above with 6 ventral midbrains pooled per sample, and sorted for CD45^+^ CD11b^+^ CX3CR1^+^ Ly6C^-^ Ly6G^-^ and NK1.1^-^ cells on a BD FACsAria. Nuclei isolation of sorted cells was performed according to manufacturer’s protocol (CG000169, 10x Genomics). Sorted nuclei were separated into single droplets or Gel beads in Emulsion (GEMs), which encapsulates each nuclei within a microgel bead containing oligonucleotides with unique barcodes and a Spacer sequence establishing barcode attachment to transposed DNA fragments to establish the ATAC library. Following GEM incubation, barcoded DNA fragments were released, amplified, and used to construct sequencing-ready ATAC libraries. Following library formation and quality control, samples were sequenced at UAB Genomics Core with an Illumina NovaSeq at an average depth of 25,000 reads per cell and a target of 7,000-15,000 nuclei per sample. Two biological replicates for each group were processed separately.

### scATAC-seq analysis

scATAC analyses followed similar standardized workflows as scRNA-seq analysis described above, using ATAC-seq specific software from Cell Ranger, Signac (Seurat), Monocle3, and Cicero. Data was aligned, filtered for profile quality, and analyzed using Signac (Seurat). Pseudotime and co-accessability analysis were conducted with Monocle3 and Cicero, respectively. TF motif enrichment analysis was performed for each cell using chromVAR.

### Single-nucleus RNA-sequencing (snRNA-seq) analysis

To evaluate the conservation of the late DAM state and its regulatory program in human PD, we utilized a previously published snRNA-seq dataset of the human substantia nigra (n = 13 PD cases and n = 15 age-matched controls), processed as previously described (Al Dalahmah et al., 2025). Following microglia subclustering, differentially expressed genes in each cluster were identified using Seurat, filtering for genes expressed in at least 25% of cells in either the cluster or remaining cells, with an average log2FC > 0.25 and an adjusted p-value < 0.05. To quantify the expression of the mouse-derived late DAM signature across human microglial states, module scoring was performed using AddModuleScore (Seurat). To model transcriptional changes associated with progression toward the late DAM state, trajectory inference was performed using Slingshot, designating the quiescent microglia as the starting state and the late DAM state as the terminal endpoint. Pseudotime values and lineage curve weights were extracted using slingPseudotime and slingCurveWeights, respectively. These matrices were subsequently used as inputs for tradeSeq to model gene expression dynamics along the inferred trajectory. Smoothed expression curves were predicted across 100 evenly spaced pseudotime points using predictSmooth. Genes exhibiting significant pseudotime-associated expression changes were identified using associationTest, followed by Benjamini–Hochberg correction. For pathway-level interpretation, genes were ranked based on the direction and significance of their pseudotime-associated expression changes, and targeted gene set enrichment analysis was performed to assess the enrichment of RelA, Fos:JunD (AP-1), and Irf4 motif-associated gene sets derived from the mouse AAV2-GFP/AAV2-SYN scATAC-seq dataset along the trajectory toward the late DAM state.

### Statistical analysis

Statistical analyses were performed using GraphPad Prism 10 software (v10.6.1). Flow cytometry experiments utilized 3 to 6 independent samples per group, with 1-4 ventral midbrains pooled per sample. Data were analyzed using unpaired two-tailed t-tests, one-way ANOVA, or two-way ANOVA, and are presented as mean ± SD or mean ± SEM, as indicated in the figure legends. Statistical significance was defined as *p ≤ 0.05, **p ≤ 0.01, ***p ≤ 0.001, ****p ≤ 0.0001. Complete statistical details and sample sizes for each group are provided in the figure legends.

## Data Availability

Source data including raw flow cytometry files, quantified data, and statistical analyses, are available in the Zenodo repository at https://doi.org/10.5281/zenodo.22864677. The single-cell ATAC-seq and single-cell RNA-seq data generated in this study will be deposited in the NCBI Gene Expression Omnibus (GEO); the accession number is pending and will be provided prior to publication. Previously published human single-nucleus RNA-seq data analyzed in Fig. 5 are available in GEO under accession number GSE253462. Further requests for data can be directed to the corresponding authors.

## Acknowledgements

The authors thank the University of Alabama at Birmingham (UAB) Flow Cytometry and Single Cell Core Facility for their assistance with single-cell experiments; Karen L. Eskow Jaunarajs and Isidoro Cobo for manuscript discussion and critical feedback; Duy Pham for guidance on figure preparation; and Josh Randolph for maintenance of all mouse lines. This study was supported by grants from the Aligning Science Across Parkinson’s (grant no. 000375), the Michael J. Fox Foundation for Parkinson’s Research, and the Parkinson Association of Alabama.

## Author Contributions

Y.Y., D.A.F., and A.S.H. designed the studies together and wrote the manuscript. Y.Y. and A.Z. were responsible for stereotaxic surgery procedures. Y.Y., A.T.M., J.M.W., A.Z., and J.R. executed IHC and flow cytometry experiments, and Y.Y. performed the corresponding analysis. D.A.F., Y.Y., and A.S.H. designed the single-cell sequencing studies, and D.A.F. and Y.Y. analyzed the data and created the corresponding figures. Y.G. and O.A. performed the analysis of the human sequencing data. Y.Y., D.A.F., and A.S.H. designed and drafted the manuscript figures. All authors read and approved the final manuscript.

## Supplementary Figure Legends

**Supplementary Fig.1. DAM accumulate near α-syn pathology and exhibit elevated cytokine and chemokine production a** Representative flow plots showing gating strategy for lymphocytes, monocytes/macrophages, and distinct microglial states. **b** Quantification of frequencies of lymphocyte and monocyte/macrophage infiltration in *Spp1*-IRES-tdTomato mice at 1 and 3 months post-AAV2-GFP or AAV2-SYN transduction. **c** Representative immunofluorescent images of Clec7a^+^ Spp1^+^ DAM around α-syn-transduced neurons in *Spp1*-IRES-tdTomato mice at 3 months post-AAV2-SYN transduction. Tissues are labeled with Spp1 (red), Clec7a (Cyan), and AAV2-SYN (green). Images are taken at 60× magnification and scale bar is 20 μm. **d** Experimental schematic of intracellular cytokine/chemokine staining following LPS and IFN-*γ* stimulation in *Spp1*-IRES-tdTomato mice at 3 months post-AAV2-SYN transduction. Microglia (Mg) were gated on live CD45^+^ CD11b^+^ Ly6C^-^ CD38^-^ P2RY12^+^ and further categorized as quiescent Mg (MHCII^-^), activated Mg (MHCII^+^), early DAM (MHCII^+^ CD11c^+^ Spp1^-^), and late DAM (MHCII^+^ CD11c^+^ Spp1^+^). **e, f** Representative flow plots showing the frequency of TNFα^+^ and IL-1β^+^ (**e**) and CCL5^+^ (**f**) cells across microglial states. **g** Quantification of (**e**) and (**f**). *n* = 4 per group, with 2 ventral midbrains pooled per *n*. One-way ANOVA, with Tukey’s test. Mean ± SD are shown. ***p ≤ 0.001, ****p ≤ 0.0001. **h** Experimental schematic of immunofluorescent staining of Clec7a in WT, MHCII knockout (KO), and CD4 KO mice at 3 months post-AAV2-SYN transduction. **f** Representative immunofluorescent images of Clec7a^+^ DAM around α-syn-transduced neurons (indicated with white arrowheads) in WT, MHCII knockout, and CD4 knockout mice.Tissues are labeled with Clec7a (red), and AAV2-SYN (green). Images are taken at 60× magnification and scale bar is 20 μm. Panel **d** created in BioRender. Yang, V. (2026) https://BioRender.com/80l5wuz

**Supplementary Fig.2. α-syn overexpression increases chromatin accessibility at DAM-associated loci a** Bar graph showing the log2 fold change of each cluster in AAV2-SYN compared to AAV2-GFP **b** scATAC-seq peaks showing chromatin accessibility at the *Gpnmb* locus. Gray shaded boxes highlight increased accessibility in clusters 11, 4, and 10.

**Supplementary Fig.3. Recombination of *TMEM119*-CreERT2 is microglia-specific a** Experimental schematic designed to assess cre recombination efficiency in *TMEM119*-CreERT2^+/-^ tdTomato^+/-^ mice. Mice were intraperitoneally injected with or without 3 mg tamoxifen per day for 5 consecutive days. 2 weeks post-tamoxifen administration, recombination efficiency were analyzed using flow cytometry. **b, c** Representative flow plots (**b)** and quantification (**c**) showing recombination efficiency across microglia, BAMs and blood monocytes. *n* = 4 per group. Two-way ANOVA, with Sidak’s test. Mean ± SD are shown. ***p ≤ 0.001, ****p ≤ 0.0001. Panel **a** created in BioRender. Yang, V. (2026) https://BioRender.com/hb3u0rw

**Supplementary Fig.4. *RelA* cKO reduces activated microglia while *Irf4* cKO suppresses DAM and proliferating microglia a** Integrated UMAP plot with cell-type labels transferred from the previous published AAV2-GFP/AAV2-SYN dataset. **b** Bar graph showing the log2 fold change of each cluster in *RelA* cKO or *Irf4* cKO, compared to Cre-ctrl

